# Inter-strain diversity among coastal picocyanobacteria across salinity gradients

**DOI:** 10.1101/2025.10.19.683324

**Authors:** M. Berthold, N.M. Omar, A. MacCormack, M. Savoie, M. Hagemann, D.A. Campbell

## Abstract

Strains of the picocyanobacteria genera *Synechococcus* and *Cyanobium* occur in cold-temperate zones from coastal waters to the open ocean, spanning a range of salinities and water depths. To thrive across different salinities, picocyanobacteria strains acclimate internal turgor by expending metabolic energy to accumulate compatible solutes and extrude inorganic ions. We grew five picocyanobacterial strains, derived from a range of source habitat salinities, under a matrix of light and salinity levels. Although Chl is a general proxy for phytoplankton biomass, Chl/cell is itself a key component of acclimation. Indeed, for all strains, except the brackish water CZS48M, the conditions supporting the highest Biovolume-specific growth rates shifted upwards towards higher salinities and PAR, compared to conditions for maximum Chl-specific growth. Therefore, the apparent optimal niche for a strain varies depending upon the metric used to track growth.

Strains CZS48M and CZS25K derived from brackish lagoons of the Baltic Sea may be true ‘brackobionts’, with maximal growth at brackish salinities and high PAR, coinciding with their maximal metabolic capacities. CZS48M and CZS25K both also show high metabolism under higher salinity and low PAR, suggesting a situation of rapid metabolism, but lower achieved growth, under stress at the edges of their environmental tolerance ranges. Conversely, NIES981 (full marine, East China Sea) shows rapid metabolism under high, but also under low, salinities. The preferred salinity of each strain is consistent with its genome-encoded capabilities to synthesize different compatible solutes. In general, across strains, maximal metabolic rate is often offset from growth rate optima, showing conversion of metabolic electrons to biomass varies widely depending upon strain and condition, likely partly reflecting the costs of turgor regulation at the limits of salinity tolerance.

## Introduction

Coastal waters worldwide contain a global brackish microbiome [1], which is in part dominated by cyanobacteria. Cyanobacteria are the only prokaryotic phylum capable of performing oxygenic photosynthesis. According to their form of RubisCO and carboxysome structure cyanobacteria are divided into alpha- and beta-cyanobacteria [2]. The alpha-cyanobacteria form an evolutionary clade, described as picocyanobacteria because of generally smaller cell sizes compared to beta-cyanobacteria. Picocyanobacteria are ubiquitous primary producers, with diverse strains exploiting different ecological niches, in spite of small genome sizes for individual strains. Strains of the picocyanobacteria genera *Synechococcus* and *Cyanobium* frequently occur in cold-temperate systems, from coastal waters to the open ocean, where they contribute substantially to annual global CO2 fixation [3]. *Synechococcus* and *Cyanobium* are divided into several clades based upon cell diameter, DNA base composition, pigment composition, and ecophysiological traits, with different claded occurring across a range of salinities and water depths [4,5] Awareness of the wide distribution of these picocyanobacteria is increasing with metagenomic approaches [6], which notably show a changing picocyanobacterial genome repertoire in the saline gradient of the brackish Baltic Sea [7]..

To thrive in ecosystems of different salinities, spanning hypersaline, marine, brackish or freshwater habitats, cyanobacteria need to acclimate internal turgor toward external salinity. Salt acclimation has been analyzed mainly among the beta-cyanobacteria, which accumulate specific compatible solutes and actively extrude inorganic ions upon salt stress [8]. Specific compatible solutes determine maximum salt tolerance; for example freshwater and terrestrial strains typically accumulate sucrose and/or trehalose, while marine strains accumulate glucosylglycerol (GG) and glucosylglycerate (GGA); whereas strains from hypersaline environments accumulate glycine betaine (GB) or homoserine betaine [9–11]. Genes for GG synthesis are frequently found in marine picocyanobacteria of the genus *Synechococcus* [12], whereas only one *Prochlorococcus* strain from the Red Sea with expressed genes for GG synthesis has been detected [13]. Laboratory analyses of a few alpha picocyanobacteria revealed that GG, GGA but also GB are synthesized upon salt stress in *Synechococcus* spp., while *Prochlorococcus* spp. responded to different salinities by accumulating sucrose, GGA and/or GB [14].

It is likely that several picocyanobacteria from one clade can form meta-populations of genetically slightly distinct strains which, through their inter-strain diversity, are able to exploit a wider range of environmental conditions, including salinities [15]. Such strain diversity of picocyanobacteria is ecologically relevant, as they may represent sentinels of a warming future [16–18], which will likely benefit smaller phytoplankton cells. Community shifts towards smaller cells have several implications for the functioning of ecosystems, including grazing evasion [19], altered sinking [20], and increased optical attenuation per unit of phytoplankton biomass [21,22]. These characteristics of small picocyanobacteria impact biogeochemical cycles and nutrient sequestration rates, other phytoplankton taxa, and ultimately the transfer of carbon through the food web. Indeed, recent studies suggest that a community shift towards dominance by small phytoplankton caused a decline in mussel size in the Baltic Sea, likely as a response to lowered energy transfer by small phytoplankton [23]. Ecophysiological characterization of different strains, coupled with analyses of their underlying genomic capacities, can thus contribute to improved predictive models of when and where such strains can be expected to rise to dominance. A problem is how to efficiently test and functionally characterize the rapidly accumulating abundance of available, identified, and even sequenced strains.

Picocyanobacterial strains have been clustered into genomic clades, primarily by using 16s rDNA [24,25], or grouped by physiological characters, such as pigment composition [4,26], both of which may fail to capture the ecological niches of strains. We herein implement a high-throughput approach to rapidly describe the growth optima and photophysiologies of picocyanobacteria, starting with a case study of five *Cyanobium* or *Synechococcus* strains, related based on conserved sequences including 16s rDNA, but originating from geographically and ecologically diverse regions of the Gulf of Mexico, Baltic Sea, Chesapeake Bay, and coastal Pacific. Our efforts are motivated by the rising omnipresence of picocyanobacteria across brackish environments [27]. We hypothesize that phenotypically similar, but genotypically and physiologically distinct strains, form meta-populations across salinity gradients. We show that strain-specific salinity optima can be differentially expressed in terms of fastest growth rates, highest quantum yield, or metabolic rate.

## Material and methods

### A. Strains, culturing, & media preparation

We chose five picocyanobacterial strains to test differential responses to salinity (parts per thousand, ppt) and Photosynthetically Active Radiation (PAR, μmol photons m^-2^ s^-1^). Two strains (CZS25K & CZS48M) were isolated from the Baltic Sea at a brackish lagoon in Germany (Darss-Zingst Lagoon System), and were previously maintained in freshwater BG11. One strain CCMP1333 (WH5701) was isolated from the Long Island Sound, USA, and selected because of close 16s rDNA homology with the two Baltic lagoon strains. CCMP1333 is also one of the earlier completely sequenced cyanobacterial genomes. Two other strains were selected because of their capacity to grow in full marine media and similar physiological characteristics or genomic similarities to the other strains. CCMP836 was isolated from the Gulf of Mexico, and chosen because of pigment characteristics and 16S rDNA sequences close to the Baltic strains [4,5] NIES-981 was isolated close to Hokkaido island, Japan, but again shows a 16s rDNA sequence similar to the Baltic strains. Strains were initially grown in fully enriched BG11 media [28,29] made with either freshwater (CZS25K and CZS48M) or full saline artificial seawater (CCMP1333, CCMP836, NIES981). Subsequently, CZS25K and CZS48M were shifted to BG11 at a salinity of 4 ppt to increase the growth rates of mother inoculant cultures. All strains were pre-cultured on a shaker, at 22 ℃, with 12:12 light:dark at 60 – 80 µmol photons m^-2^ s^-1^ from white fluorescent bulbs. Pre-cultures were diluted 1 to 4 in fresh media every seven days to keep them actively growing. We chose BG11 media as it contains sufficient Nitrogen (N) in the form of nitrate to keep batch cultures nutrient replete, although using nitrate instead of ammonium as a primary N source impacts the growth and physiology of the cells [30]. BG11 media with different salinities were created using artificial sea water mixed with deionized water (Milli-Q) to salinity levels of 1, 4, 11, 18, 25 and 32 ppt. The media tends to flocculate with increasing salinities due to increasing pH. Thus, we titrated each media back to the recommended pH of 7.5 using 1 M hydrochloric acid, upon which flocculation and precipitates re-dissolved. We did not tightly control the media dissolved inorganic carbon (DIC), but the BG11 was prepared with an initial concentration of 0.189 mM NaHCO_3_. The atmospheric CO_2_ in the culture chambers was approximately 750 ppm, leading to an estimated initial equilibrium [DIC] at pH 7.5 of ∼1356 mM [31].

### B. Experimental design; absorbance and fluorescence measurement regimes and data handling

We used 24-well plates (Falcon, 2.5 ml volume per well, No. 351147) to culture each strain at six different salinity levels (six columns) with four replicates (four rows). Each plate run was then replicated three times within 8 weeks, making a total of three separate plate replicates with four technical well replicates per each combination of strain, light level, and salinity level. Three plates for one strain were simultaneously cultured at 30, 100, and 300 µmol photons m^-2^ s^-1^ in a common growth chamber for consistency across different growth PAR. Wells held one magnetizable micro-stir bar (V&P Scientific Inc.). Each day cells in well plates were resuspended using an external magnet (V&P Scientific Inc.) and then immediately measured in a CLARIOstar (BMG LABTECH) in top read mode, with an excitation at 440 nm for a fluorescence emission spectral scan from 500 to 750 nm, and an absorbance spectral scan, 400 to 750 nm. BG11 media with different salinities had different inherent OD values. We read plates with media alone before every inoculation to correct for well and media-specific absorption and thus blank-correct each culture across the spectral scans. Furthermore, batches of BG11 media had slightly different salinities, so each batch was assigned a unique ID to track final salinity in each culture well.

For import, tidying, analyses and presentation of data we used R [32] running under RStudio [33] and packages [34–41].

### C. Cell count & chlorophyll calibrations

From each plate, after each doubling of Chlorophyll a (Chl) content, tracked as OD_680nm_ – OD_750nm_, plates were opened in a culture hood and a culture from each replicate row of wells, representing all six salinity levels, were harvested for further analyses. These samples after each doubling were analyzed for Chl, cell number, and their photophysiological characteristics. For Chl *a*, 20 µl of culture were mixed with 2 ml of 90 % acetone:DMSO (3:2 ratio), incubated for 30 min in the dark and analyzed with a Turner Trilogy (Turner Designs, San Jose, CA, USA) fluorometer, calibrated using Chl *a* standards, with relative fluorescence units converted to Chl *a* in µg mL^-1^. These Chl *a* concentrations were compared to respective (OD_680nm_ – OD_750nm_) values from the same sample and day to generate strain- and growth light-specific regression models for converting daily (OD_680nm_ – OD_750nm_) to absolute [Chl a] (S1 Table). Chl content is a component of photoacclimation, so in parallel we counted cells from 10 µl samples of culture using a hemocytometer and a Leica microscope at 400x magnification. Those cell counts were compared to the respective OD_750nm_ values from the same sample and day to generate strain- and growth light-specific regression models for converting daily OD_750nm_ to cells mL^-1^ (S2 Table). Cell biovolumes varied across the strains and growth conditions, so we estimated total biovolume trajectories by multiplying cell counts by strain- and condition-specific biovolumes. We were thus able to calculate Chl- and biovolume-specific growth rates for each strain.

### D. Growth analyses

The daily well specific [42] measures of [Chl *a*], derived from strain- and light-specific regressions of (OD_680nm_ – OD_750nm_), and Cell Counts, from regressions of OD_750nm_, were fit using a modified Gompertz model, chosen to account for any lag-phase. The modified Gompertz model as described by Zwietering et al. [43] is given as:

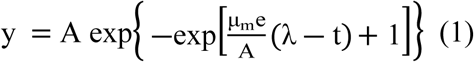

where A is the maximum population size [A = ln(N_∞_/N_0_)], λ is the lag time, and μ_m_ is the maximum specific growth rate.

Modified Gompertz growth curves were fitted using non-linear model fitting functions [44]. Growth models were fit separately for each strain, salinity and plate replicate (n = 3) based on the pooled four (start of experiment) to two (end of experiment) technical replicate wells per plate for each salinity (S3 Table).

### E. Photophysiology

The remaining 2 mL per well harvest was used for photophysiological characterizations of each culture, at every salinity and light level, repeatedly across the growth trajectories. We used Fast Repetition Rate fluorometry [45] (FRRf, Solisense, USA) to apply series of flashlets to drive induction/relaxation trajectories. We used a double tap protocol [46], where an FRRf induction/relaxation trajectory is applied on top of an actinic light level, immediately followed by another induction/relaxation after 1 s in the dark, to allow re-opening of PSII. Actinic light steps of 10 s were applied at 0, 20, 40, 80, 160, and 320 µmol photons m^-2^ s^-1^, at 445 or 590 nm. FRRf excitation flashlets were applied at the same waveband, 445 or 590 nm, as the actinic light steps.

We used the onboard Solisense LIFT software to fit an induction/relaxation model [45] for each FRRf induction/relaxation trajectory. From the model fits of the inductions, we took the maximum quantum yield of Photosystem II (PSII) (F_V_/F_M_) as the photochemical capacity of the cultures. The relaxations were fit with a three-component exponential decay model, generating lifetimes τ_1_, τ_2_, and a slow τ_3_ of small amplitude. We calculated an amplitude weighted average of τ_1_ and τ_2_ for the re-opening of PSII by downstream electron transport as a proxy for down stream metabolic capacity to use electrons.

### F. Genomic analyses

Predicted protein sequences derived from the genomes of *Cyanobium* spp. (strains CZS25K, CZS48M, and NIES981) and *Synechococcus* spp. (strains CCMP836 and WH5701) were obtained from NCBI [47,48]. We used gene sequence probes (S4 Table) to search the genomes for the presence of known genes encoding proteins involved in osmolyte production and ion transport, using BLAST+ v2.13.0 [49] with an e-value of 1*10^-10^ as a cutoff for genes homologous to the probe sequences. Counts of detected hits were pooled based on gene name and used in subsequent analyses.

### G. Generalized additive modelling

Estimated exponential growth rates specific for [Chl *a*] and for biovolume were used to generate generalized additive models [50,51] for growth of each strain, with salinity and light level as explanatory variables (S5 and S6 Tables) . Predicted growth rate values from the GAM models were filtered out if the standard error on the estimate was greater than 15% of the fitted value. We were thus able to generate modelled response surfaces of growth rates for each strain across the matrix of tested salinity and light growth conditions. We present strain-specific GAM estimates for growth optima by plotting the top 10% fastest predicted growth rates for strain, vs. salinity and light. In parallel, we plotted the strain which had the highest GAM estimated growth rate under a given combination of salinity and light, for a ‘winner takes it all’ approach to define which strain would potentially dominate under given conditions.

In parallel, we fit GAM of F_V_/F_M_ (S7 Table) as a metric of photochemical potential across the matrix of conditions. For metabolic capacity across conditions, we took the weighted-average τ_PSII_ from the amplitudes and duration of the τ_1_ and τ_2_ extracted from the fits of the re-opening of PSII after closure, taken after 1 s of darkness, immediately after actinic light steps closest to the respective growth light, for each salinity condition and strain. For better conceptual comparability with growth rates, for fitting GAM (S8 Table) we used the reciprocal 1/τ_PSII_ which is the rate constant (s^-1^) for electrons transported away from PSII. Again, for presentation of the GAM model for optimal conditions for each strain, we chose the upper 10% of F_V_/F_M_ or 1/τ_PSII_ across growth conditions and strain. Biovolume-specific growth rates were compared to F_V_/F_M_ and 1/τ_PSII_, to uncover patterns in the conversion of photochemistry and metabolic electrons to biomass across salinity and PAR (S9 Table).

## Results

### i) Salinity-light niches for chlorophyll- and biovolume-specific growth rates across strains

We summarized growth rates for the strains across the matrix of PAR and salinity conditions using GAM (Fig 1 and Fig 2). To identify the optimal growth conditions for each strain, we plot the regions where each given strain achieved the top 10% of their Chl-specific, or Biovolume -specific, growth rates (Fig 1).

**Fig 1.**
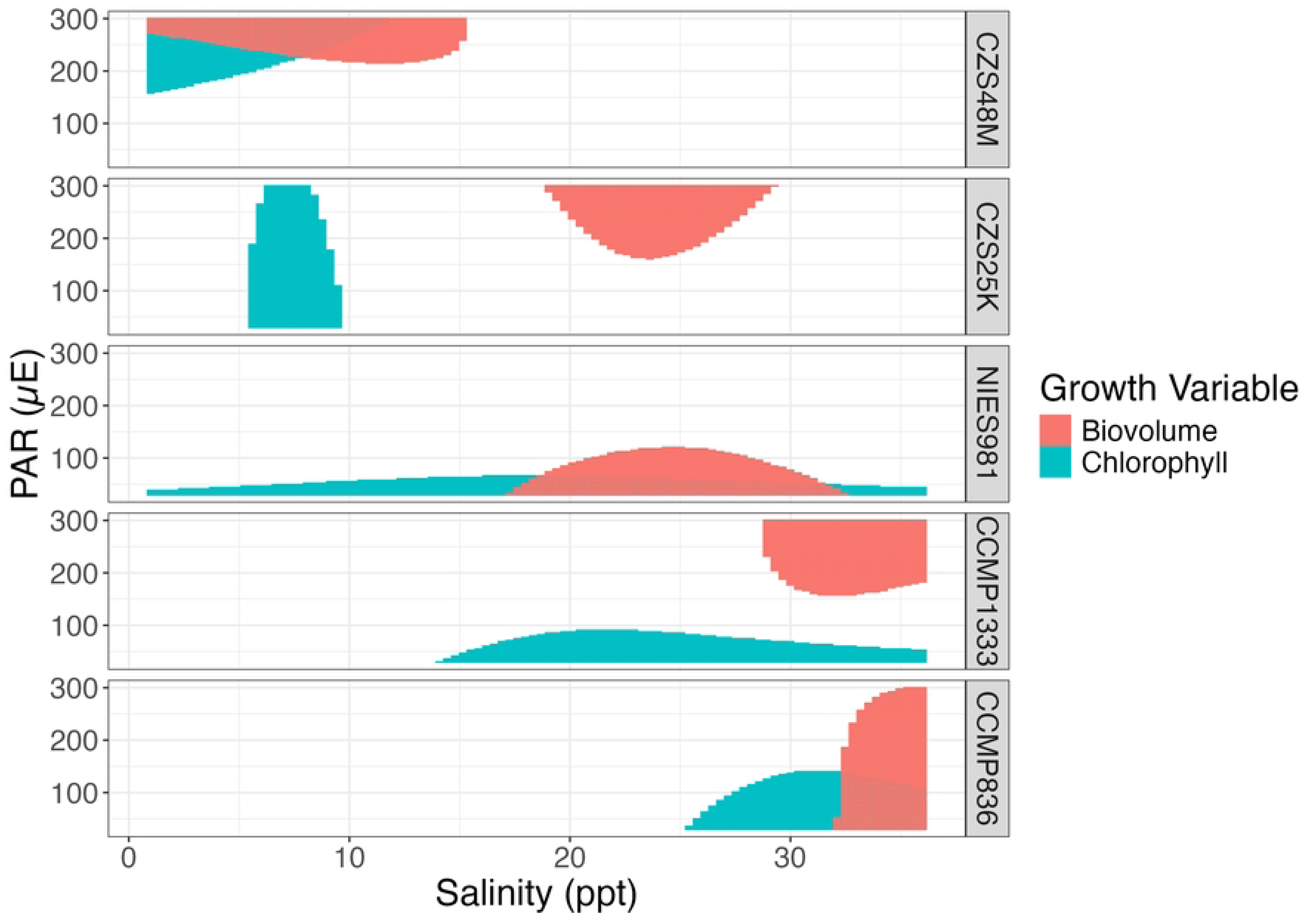
Comparison of Conditions supporting Maximum Biovolume-Specific or Maximum Chlorophyll-Specific growth rates (d^-1^) of picocyanobacterial strains, growing across a matrix of salinities (ppt) and light levels (PAR, µmol photons m^-2^ s^-1^, µE). Each panel shows growth results from a different strain, ordered top to bottom, from low to high salinity tolerance. Red regions show the conditions where a strain achieved the top 10% of its Biovolume-specific growth rates, while green regions show the conditions where a strain achieved the top 10% of its Chl-specific growth rates. The growth rates used for Generalized Additive Modelling were taken from Gompertz fits of changes in total biovolume (µm^3^ L^-1^) or total Chl (µg L^-1^) over time (d^-1^) under the respective PAR and salinities for each strain (S3 Fig).

Chl-specific growth rates of CZS48M (brackish, Baltic Sea) were highest at salinities of 0 – 15 ppt, and at high light of 220-300 µmol photons m^-2^ s^-1^. Similarly, the Biovolume-specific growth rates of CZS48M were highest around salinities of 0 – 15 ppt and again at higher PAR of 175 – 300 µmol photons m^-2^ s^-1^, in contrast to all other strains that showed offsets in the light levels for maximal Chl *a* vs. biovolume growth rates (Fig 1).

Chl-specific growth rates of CZS25K (brackish, Baltic Sea) were highest around salinities of 5 – 10 ppt, extending across the tested light gradient of 30 – 300 µmol photons m^-2^ s^-1^. Interestingly, the highest Biovolume-specific growth rates of CZS25K were shifted upwards to salinities of 20 – 25 ppt and higher PAR of 175 – 300 µmol photons m^-2^ s^-1^, showing an acclimatory offset in conditions favouring Biovolume-vs. Chl-specific growth rates (Fig 1).

Chl-specific growth rates of NIES981 (full marine, East China Sea) were achieved across salinities of 0 – 32 ppt, but only at low light of 30 µmol photons m^-2^ s^-1^; in contrast Biovolume-specific growth rates for NIES981 were highest around salinity of 25 ppt but extended up to higher PAR of 100 µmol photons m^-2^ s^-1^ (Fig 1).

Chl-specific growth rates of CCMP1333 (full marine, Long Island Sound) were highest around salinities of 15 – 35 ppt, at low light of 30 – 100 µmol photons m^-2^ s^-1^, but the highest Biovolume-specific growth rates were offset to salinities around 30 – 35 ppt and higher light of 150 – 300 µmol photons m^-2^ s^-1^ (Fig 1).

Chl-specific growth rates of CCMP836 (full marine, Gulf of Mexico) were highest around salinities of 25 – 35 ppt, at PAR of 30 – 100 µmol photons m^-2^ s^-1^ (Fig 1) while the highest Biovolume-specific growth rates were achieved above salinity of 32 ppt, across the measured PAR from 30 – 100 µmol photons m^-2^ s^-1^ (Fig 1). Thus, for CCMP836, conditions for the highest Biovolume-specific growth shift upwards to higher salinities and light, compared to Chl-specific growth.

**Fig 2.**
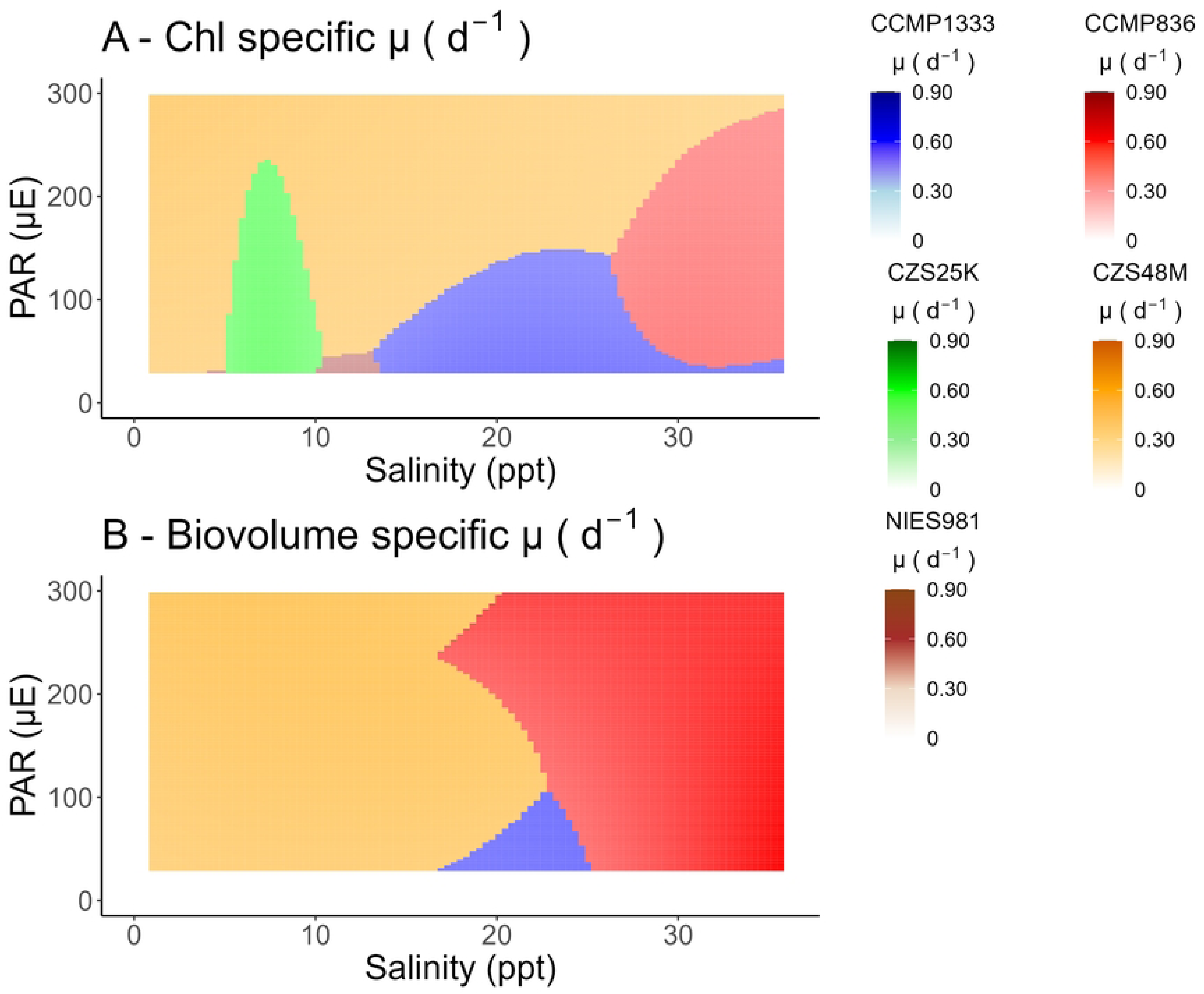
Comparisons of Chlorophyll- or Biovolume-specific growth rates (d^-1^) of picocyanobacterial strains growing across a matrix of salinities (ppt) and light levels (PAR, µmol photons m^-2^ s^-1^). For each combination of light level (PAR, µmol photons m^-2^ s^-1^) and salinity (ppt), the strain (colour), cultured separately, achieving the highest Chl-specific growth rate (d^-1^) or biovolume-specific growth rate, is indicated by shading, in a ‘winner takes it all’ comparison of potential competitive advantage among strains across conditions. The growth rates used for Generalized Additive Modelling were taken from Gompertz fits of changes in total biovolume (µm^3^ L^-1^) or total Chl (µg L^-1^) over time (d^-1^) under the respective PAR and salinities for each strain.

As an alternative to strain-specific maximum growth rates (Fig 1), Fig 2 instead compares which strain achieves the highest Chl-specific or Biovolume-specific growth rate under a given combination of PAR and salinity.

CZS48M (brackish, Baltic Sea) is a Chl-specific growth winner across the range of salinities under high PAR, but is outcompeted by other strains at lower PAR, particularly as salinity increases. In contrast, CZS48M only achieves the highest Biovolume-specific growth under salinities less than ∼ 20 ppt, but across the full range of PAR (Fig 2).

CZS25K (brackish, Baltic Sea) is only a winner for Chl-specific growth at salinities around 8 ppt, up to ∼ 250 µmol photons m^-2^ s^-1^, but never achieves the highest Biovolume-specific growth rates among strains (Fig 2).

NIES981 (full marine, East China Sea) achieves the highest Chl-specific growth in a small region of moderate (∼ 10 ppt) salinity and PAR below 30 µmol photons m^-2^ s^-1^, but never achieves the highest Biovolume-specific growth rate among strains (Fig 2).

CCMP1333 (full marine, Long Island Sound) achieves the highest Chl-specific growth at salinities ∼15 -25 ppt, and PAR below 150 µmol photons m^-2^ s^-1^ (Fig 2). CCMP1333 achieves the highest Biovolume-specific growth rates in a smaller region centred at salinity of ∼ 22 ppt, below 100 µmol photons m^-2^ s^-1^ (Fig 2).

CCMP836 (full marine, Gulf of Mexico) achieved the highest Chl-specific growth above salinities of ∼30 ppt, almost across the range of PAR. Similarly, CCMP836 achieved the highest Biovolume-specific growth rates across a wider range of salinities above ∼ 20 ppt, again, from low to high PAR (Fig 2).

### ii) Interactions of photochemistry and metabolism, across salinity and light ranges

**Fig 3.**
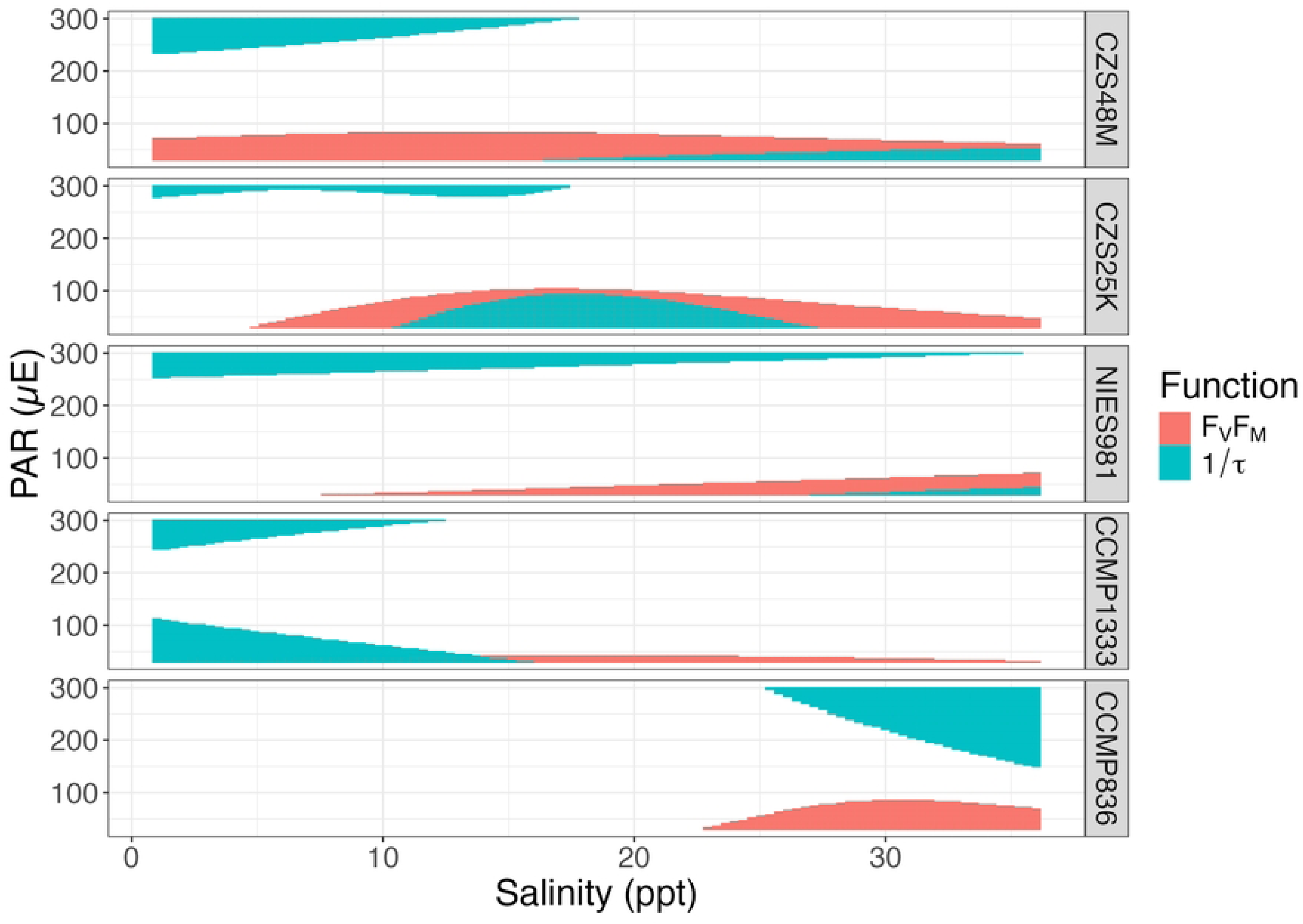
Comparison of Conditions supporting Maximum PSII Photochemistry (F_V_/F_M_) or downstream electron transport rates (1/τ) (s^-1^) of picocyanobacterial strains, growing across a matrix of salinities (ppt) and light levels (PAR, µmol photons m^-2^ s^-1^, (µE)).

Each panel shows growth results from a different strain, ordered top to bottom, from low to high salinity tolerance. Red regions show the combinations of conditions where a strain achieved the top 10% of its F_V_/F_M_ ; while green regions show the conditions where a strain achieved the top 10% of its (1/τ) (s^-1^). F_V_/F_M_ and 1/τ used for Generalized Additive Modelling were taken from fits of Fast Repetition and Relaxation fluorescence induction curves measured under the respective combinations of PAR and salinity for each strain (Fig 3).

F_V_/F_M_ values for CZS48M (brackish, Baltic Sea) were highest across the salinity range from 0 – 35, but under PAR below 100 µmol photons m^-2^ s^-1^. In contrast, CZS48M achieved maximum 1/τ under either high PAR and low salinities, or low PAR and high salinities (Fig 3).

F_V_/F_M_ values for CZS25K (brackish, Baltic Sea) were highest around 10 – 25 ppt, again under PAR below 100 µmol photons m^-2^ s^-1^. CZS25K again achieved maximum 1/τ under either high PAR and low salinities, or low PAR and intermediate salinities (Fig 3).

F_V_/F_M_ values for NIES-981 (full marine, East China Sea) were achieved at salinities above 10 ppt, but again only at low PAR of 30 µmol photons m^-2^ s^-1^. NIES-981 achieved maximum 1/τ under either high PAR and low salinities, or low PAR and high salinities (Fig 3).

F_V_/F_M_ values for CCMP1333 (full marine, Long Island Sound) were highest around a salinity of 20 ppt, under low PAR of 30 µmol photons m^-2^ s^-1^. CCMP1333 achieved maximum 1/τ under salinities below 10 ppt, under both low and high PAR (Fig 3).

F_V_/F_M_ values for CCMP836 (full marine, Gulf of Mexico) were highest around salinities of 25 – 35 ppt, at PAR below 100 µmol photons m^-2^ s^-1^, while the maximum 1/ τ was above salinities of 30 ppt and PAR above 150 µmol photons m^-2^ s^-1^ (Fig 3).

**Fig 4.**
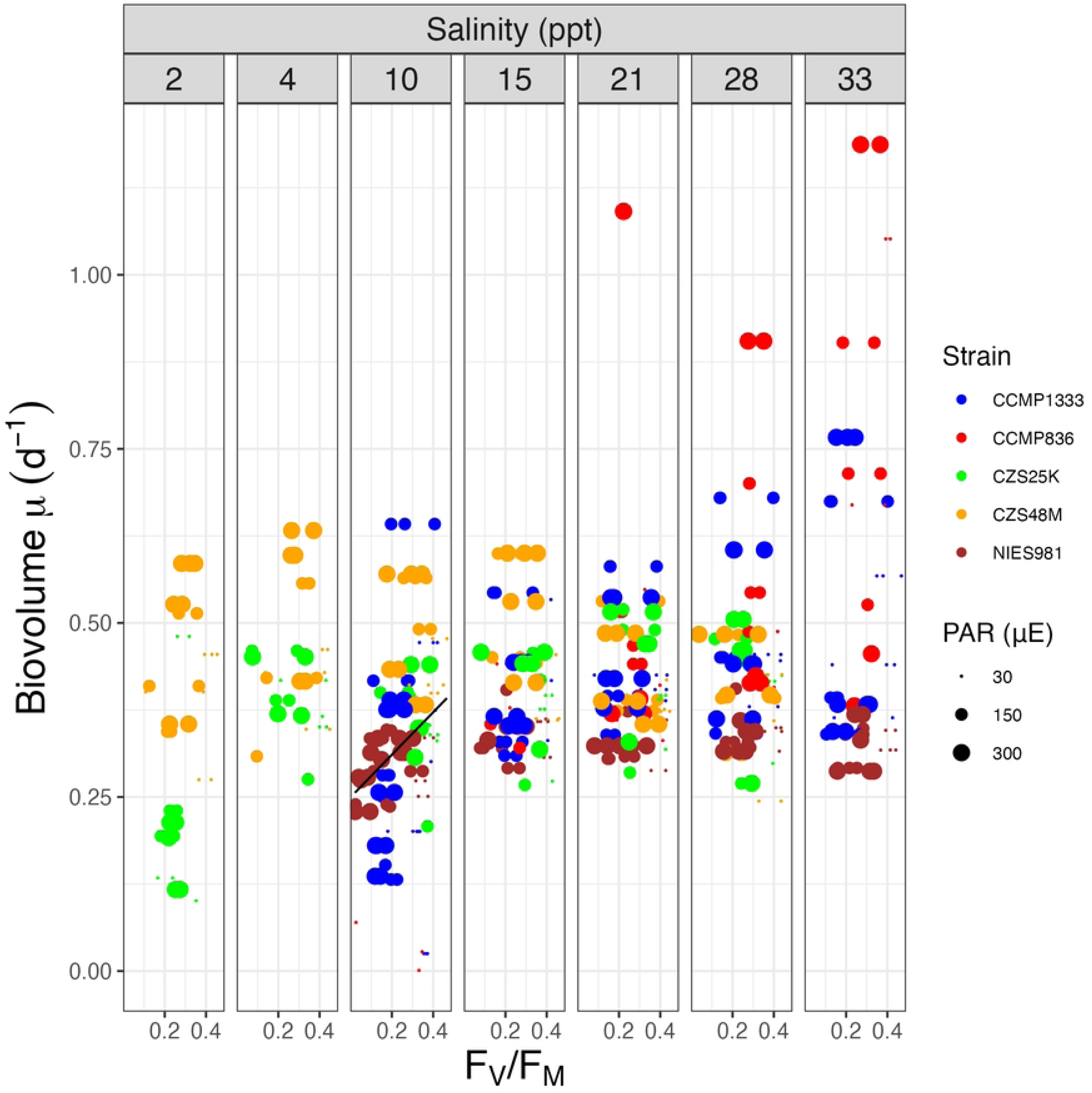
Biovolume-specific growth rates (d^-1^) of picocyanobacterial strains (symbol colours) vs. F_V_/F_M_ across a matrix of growth media salinity (ppt; columns) and growth light level (PAR, µmol photons m^-2^ s^-1^; symbol size).

**Fig 5.**
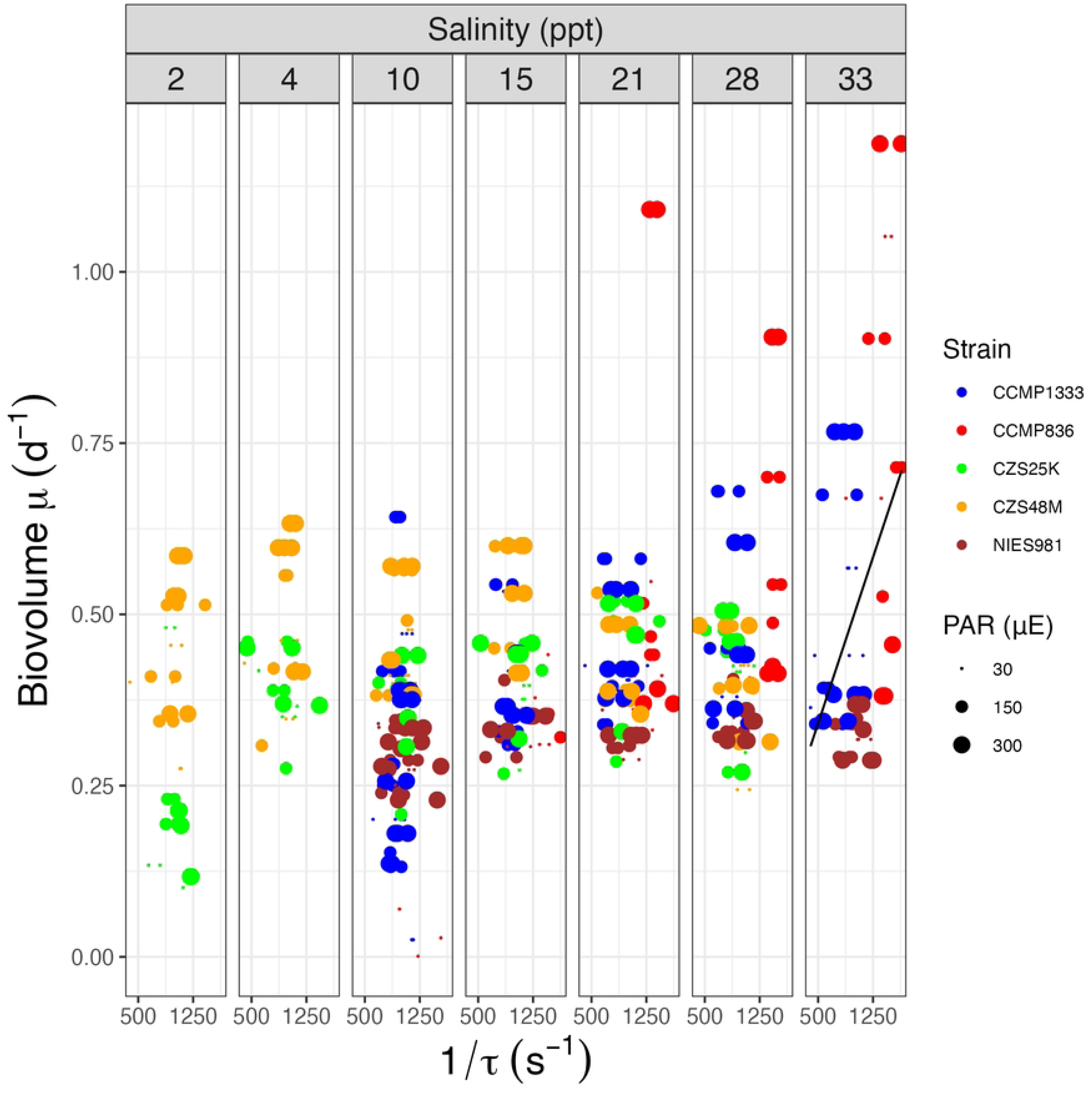
Biovolume-specific growth rates (d^-1^) of picocyanobacteria strains (symbol colours) vs. 1/τ (s^-1^) across a matrix of growth media salinity (ppt; columns) and growth light level (PAR, µmol photons m^-2^ s^-1^; symbol size).

To study if photochemical capacity (F_V_/F_M_) (Fig 4) or metabolic capacity (1/tau (s^-1^) (Fig 5) explained patterns of Biovolume-specific growth across salinities, we ran linear regression analyses (S9 Table) with final media salinity as a co-variant, while pooling data across strains and growth PAR, after preliminary analyses including these as co-variants. Since chlorophyll is itself a key component of phytoplankton acclimation, we focused on Biovolume-specific growth rates. Interestingly growth PAR from 30 to 300 µmol photons m^-2^ s^-1^ had little influence on the achieved Biovolume-specific growth rates, while, as seen in the GAM, CCMP836 and CCMP1333 stand out as achieving the highest Biovolume-specific growth rates, along with high F_V_/F_M_ (Fig 4) and 1/tau (s^-1^) (Fig 5) when growing under full marine salinity. Across strains, F_V_/F_M_ was the only significant positive predictor of growth rate under moderate salinity of ∼ 10 ppt, but not under full marine salinity (Fig 4). Conversely, 1/tau (s^-1^) was a significant positive predictor of growth only under full marine salinity, driven largely because of rapid growth and high 1/tau (s^-1^) of CCMP836 and CCMP1333 under marine salinity of ∼33 ppt (Fig 5).

### iv) Genomic analyses of salinity tolerance marker genes

**Fig 6.**
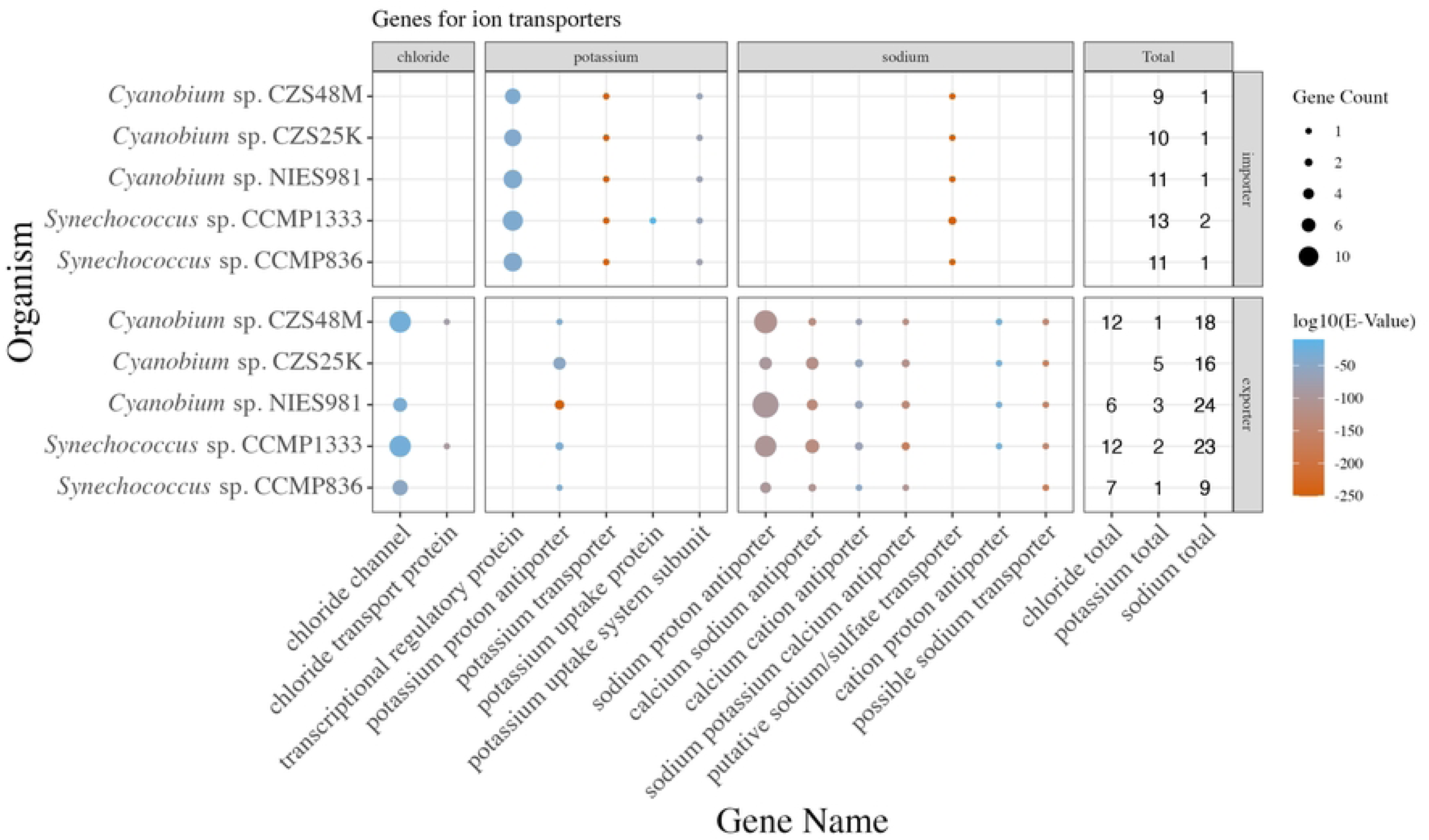
Summary of genes encoding ion transporters in study picocyanobacteria, faceted by their respective ion substrates and whether they import or export ions. Strains are ordered from lower (top) to higher (bottom) salinity tolerance. Symbol size represents counts of genes encoding each ion transporter in each genome. Colour represents log10 of the average E-value of the BLAST match between the probe sequence and the target sequence in the cyanobacterial genome. Symbol absence means no sequences encoding a given function were detected in a given genome.

Our results show differences in the growth and photosynthetic capacities among the strains, isolated from different saline environments. The original habitat of isolation, however, does not necessarily determine salt tolerance, as observed for the model cyanobacterium *Synechocystis* sp. PCC 6803, isolated from freshwater, but which can resist up to twofold seawater salinity. Therefore, we checked the genome sequences for genes encoding proteins potentially involved in salt acclimation using sequence probes derived from verified salt resistance proteins from different cyanobacteria and heterotrophic bacteria (see S4 and S10 Tables). All the investigated cyanobacteria use the ‘salt out’ strategy under saline conditions, through export of inorganic ions such as Na^+^ and C^l-^, while also accumulating compatible solutes. Our analyses of a panel of genes encoding ion transporters shows limited evidence for a shift from the brackish to the full marine strains (Fig 6). All strains possess transporters that can potentially export Na+ and Cl-out of the cell, and several transporters involved in K+ uptake, along with a single putative importer for Na+ uptake. This finding is similar to most other cyanobacteria, with no clear differences in the genetic capabilities for ion transport found among strains of different salt tolerances [8].

**Fig 7.**
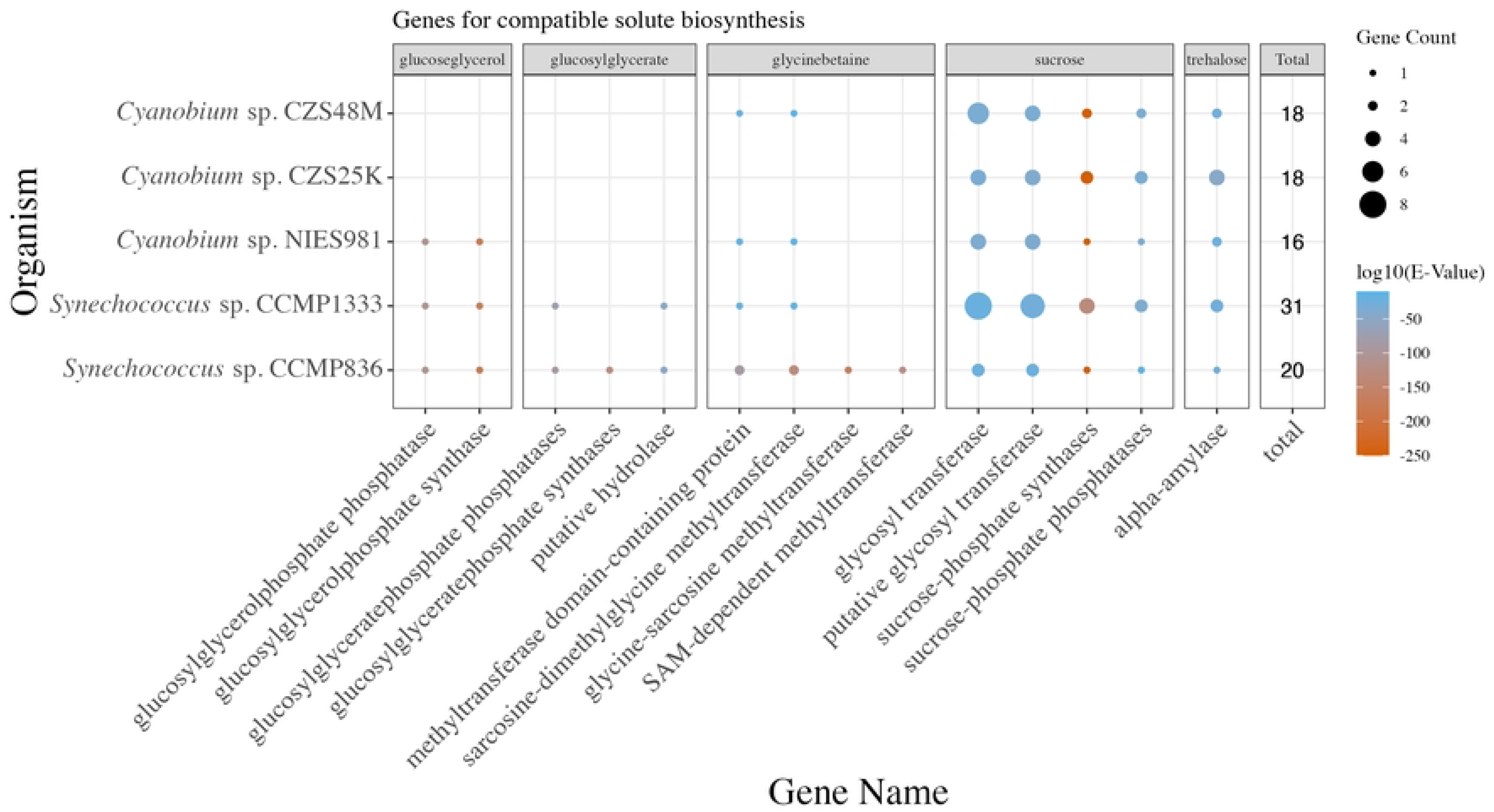
Summary of genes encoding compatible solute biosynthesis in Cyanobacteria, faceted by solute products. Strains are ordered from lower (top) to higher (bottom) salinity tolerance. Symbol size represents counts of genes encoding each ion transporter in each genome. Color represents log10 of the average E-value of the BLAST match between the probe sequence and the target sequence in the cyanobacterial genome. Symbol absence means no sequences encoding a given function were detected in a given genome.

Another picture emerged when we searched for genes encoding proteins potentially involved in the biosynthesis of different compatible solutes (Fig 7). The full marine strain CCMP836 shows the most complete panel of genes encoding proteins synthesizing a full range of compatible solutes. CCMP836 is equipped with genes for the synthesis of GG, glucosylglycerate, sucrose, trehalose, and apparently also has capacity for glycine betaine synthesis. Interestingly CCMP836 has a smaller total count of genes encoding both ion transporters (Fig 6) and compatible solutes (Fig 7), compared to the strains NIES981 and CCMP836. Both the strains NIES981 and CCMP836 show a modest drop in the count of genes encoding glycosyl-transferases and sucrose metabolism, suggestive of a shift away from carbohydrate compatible solutes towards glycine betaine (Fig 7). In contrast, the brackish water strains CZM48M and CZS25K can apparently not synthesize GG, because the GG-phosphate synthase is missing from their genomes. In all strains, genes for the synthesis of sucrose [52] and trehalose synthesis are present, which permit the brackish water strains CZM48M and CZS25K to resist moderate salinities up to 30 ppt.

## Discussion

For all strains, except the brackish CSZ48M, the conditions supporting the highest Biovolume-specific growth rates shift upwards towards higher salinities and PAR, compared to Chl-specific growth (Fig 1). Although Chl is a general proxy for biomass in phytoplankton biology, Chl/cell is a key component of acclimation [53], including responses to salinity in some phytoplankters [54]. Therefore, the apparent optimal niche for a strain will vary depending upon the metric used to track growth.

F_V_/F_M_, our proxy for photochemical potential, as expected, was maximal for all strains under low PAR, but showed a shift towards higher salinities for the full marine strains (Fig 3). For all strains except CCMP836 (full marine, Gulf of Mexico) 1/τ_PSII_, our proxy for metabolic capacity, showed a disjunct distribution of maximum values, achieved at both high, and low, PAR (Fig 3).

Each strain showed a distinct pattern of 1/τ_PSII_, across the six salinity levels and three PAR (Fig 3). In general, CCMP836 showed high 1/τ_PSII_ compared to other strains under all conditions except low salinity (Fig 3). 1/τ_PSII_ of CCMP836 (full marine, Gulf of Mexico) were highest above 30 ppt at light of 200 – 300 µmol photons m^-2^ s^-1^ (Fig 3) and were thus achieved under conditions overlapping its highest Biovolume-specific growth rates (Fig 1), with this strain showing a strong coupling between potential to generate metabolic electrons and achieved growth rate. Similarly, NIES981 (full marine, East China Sea) achieved high 1/τ_PSII_ above 30 ppt (Fig 3), slightly offset from its conditions for maximum Biovolume-specific growth rates that peaked around 25 ppt (Fig 1), but NIES981 also shows high 1/τ_PSII_ under low salinities where its growth is slower. 1/τ_PSII_ for CCMP1333 (full marine, Long Island Sound) were, surprisingly, highest at low salinity (Fig 3), below its conditions for maximum Biovolume-specific growth rates (Fig 1). 1/τ_PSII_ for CZS25K (brackish, Baltic Sea) were highest around 10 – 18 ppt and low PAR (Fig 3), overlapping the salinity range, but not the higher PAR where it achieved maximal growth rates (Fig 1). 1/τ_PSII_ for CZS48M (brackish, Baltic Sea) was highest around 10 ppt at light of 300 µmol photons m^-2^ s^-1^ (Fig 3), overlapping with its conditions for maximal growth rates (Fig 1), but 1/τ_PSII_ was again high for CZS48M at salinity of 30 and low PAR. 1/τ_PSII_ maxima are thus often offset from growth rate optima, consistent with varying ratios for conversion of metabolic electrons to biomass, depending upon strain and condition, as indeed found in wider field studies [55].

The data on growth and photosynthesis show the expected correlation between salinity and physiological performance, considering the different habitats for the strains under investigation. The better performance of marine strains at high salinities, compared to the brackish water strains, is also consistent with their genetic capacities. As expected [8] the picocyanobacteria showed no clear differences in their genetic capabilities for ion transport based upon the salinities of the originating habitats (Fig 6). In contrast their genomic complements for genes encoding proteins involved in the biosynthesis of different compatible solutes did vary systematically, with the full marine strain CCMP836 showing the most complete panel of genes encoding compatible solute synthesis whereas the brackish water strains CZM48M and CZS25K lack the gene encoding GG-phosphate synthase (Fig 7). Hence the different growth and physiological responses towards increasing salinities are explicable mainly through the different acquisition or losses of genes to synthesize appropriate compatible solutes..

Optical attenuation in the water column is another metric of competitive advantage among strains, since the success of picocyanobacteria in lagoon environments may relate to their high achieved optical attenuation relative to biomass and pigment content [56]. Growth of optical attenuation, a close correlate of Chl-specific growth rates, might therefore allow a picocyanobacteria community to reach and then maintain dominance through shading of other phytoplankton and macrophytes, particularly since the brackish cyanobacteria show only limited growth rate responses to changing PAR.

We find, based upon maximal growth rates, that CZS48M and CZS25K (brackish, Baltic Sea) may be true ‘brackobionts’, optimized for growth at brackish salinities and high PAR. Both of these strains show regions of maximal 1/τ_PSII_ coinciding with their conditions for maximal growth rates, but both also show high 1/τ_PSII_ under higher salinity and low PAR, suggesting a situation of rapid metabolism, but lower achieved growth, under stress at the edges of their environmental tolerance ranges. Conversely, NIES981 (full marine, East China Sea) shows high 1/τ_PSII_ under high salinities, but also under low salinities, again suggesting conditions of rapid metabolism under stress at the lower end of its salinity tolerance. Thus, overall, 1/τ_PSII_ shows suggestive interactions with patterns of growth rates but is a poor generic predictor of achieved growth under stressed conditions.

## Supplemental material

**S1 Table.** Strain specific estimates to convert **OD_680_-OD_750_ to μg chl mL^-1^** determined with a Turner fluorometer.

**S2 Table.** Strain-specific Biovolume estimates (μm^3^) from microscopy.

**S3 Table.** Gompertz Growth models were fit separately for each strain, salinity and plate replicate (n = 3) based on the pooled four (start of experiment) to two (end of experiment) technical replicate wells per plate for each salinity.

**S4 Table.** Genes probes used to search genomes

**S5 Table.** Generalized additive models for chlorophyll specific growth of each strain, with salinity and light level as explanatory variables

**S6 Table.** Generalized additive models for biovolume specific growth of each strain, with salinity and light level as explanatory variables

**S7 Table.** Generalized additive models for photophysiological parameters F_V_/F_M_ and metabolic capacity 1/tau (s^-1^) for each strain, with salinity and light level as explanatory variables.

**S8 Table.** Generalized additive models for photophysiological parameters F_V_/F_M_ and metabolic capacity 1/tau (s^-1^) for each strain, with salinity and light level as explanatory variables.

(TauAv_Gam.csv)

**S9 Table. Linear regression results for Biovolume-specific growth rates (d^-1^) with respect to F_V_/F_M_ or 1/tau (s^-1^)**, with covariant growth media salinity (ppt), data pooled across strains and growth lights.

**S10 Table.** Summary of genes in studied picocyanobacteria, encoding proteins potentially involved in salt acclimation, detected using sequence probes derived from verified salt resistance proteins from different cyanobacteria and heterotrophic bacteria.

## Notes

### Competing Interest Statement

The authors have declared no competing interest.

